# Ramelteon facilitates hippocampal ripple occurrence and amplitude in mice

**DOI:** 10.64898/2026.05.08.723673

**Authors:** Mahiro Nakashima, Miyuki Miyano, Honoka Kuroyanagi, Anna Sasahara, Yuji Ikegaya, Nobuyoshi Matsumoto

## Abstract

The hippocampus is essential for memory consolidation, a process mediated by high-frequency oscillations known as ripples during non-rapid eye movement (NREM) sleep. Ramelteon, a selective MT1/MT2 receptor agonist, has been reported to possess cognitive-enhancing properties; however, its impact on the fine-scale dynamics of hippocampal ripples remains unclear. We performed chronic local field potential recordings from the dorsal hippocampus and prefrontal cortex in mice. Following the intraperitoneal administration of either vehicle or ramelteon, we evaluated sleep architecture and characterized ripple properties, including occurrence rate, amplitude, instantaneous frequency, and duration during NREM sleep. Ramelteon administration significantly increased NREM sleep occupancy. Notably, we found that ramelteon significantly enhanced both the occurrence rate and amplitude of hippocampal ripples compared to the control group. While a slight increase in intra-ripple frequency was observed, other structural features, such as ripple duration and asymmetry index, remained unaffected. Our findings demonstrate that ramelteon facilitates hippocampal ripple dynamics by increasing their occurrence and synchrony during NREM sleep. Given the critical role of ripples in memory consolidation, these neurophysiological changes may underlie the procognitive effects of ramelteon.

**Graphical abstract:** 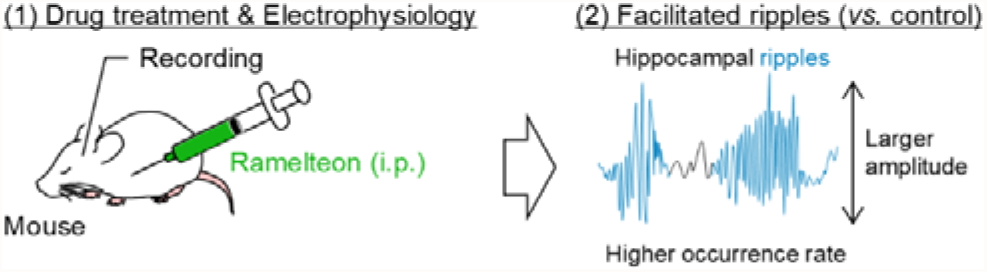

## Introduction

The hippocampal formation is a critical structure for the consolidation of episodic and spatial memories, processes mediated by coordinated neuronal activity during specific behavioral states.^2,4–6)^ During non-rapid eye movement (NREM) sleep and quiet immobility, the hippocampus exhibits transient, high-frequency oscillations (120–200 Hz) known as ripples.^7,17)^ These ripples represent the highly synchronized firing of hippocampal pyramidal cell ensembles and are often accompanied by the replay of neural sequences experienced during prior wakefulness.^1,3,5,6,8)^ Because the pharmacological or optogenetic suppression of ripples impairs memory performance, ripples are considered a physiological hallmark of memory consolidation.^6,15,21,23)^ Thus, pharmacological agents modulating ripple dynamics should be a promising strategy for treating cognitive decline and sleep-related memory disorders.

Ramelteon, an agonist of melatonin MT1 and MT2 receptors, is clinically utilized for the treatment of insomnia,^11,18,19,22)^ and may possess neuroprotective and cognitive-enhancing properties in various animal models.^9,10,24)^ Consistently, we have demonstrated that ramelteon coordinates neural oscillations in the hippocampus and prefrontal cortex (PFC) during novel object recognition.^9)^ However, it still remains unknown whether and how ramelteon affects the fine-scale properties of hippocampal ripples. Here, we chronically recorded local field potentials (LFPs) from the hippocampus and PFC in mice during sleep in the home cage and investigated how ramelteon affects ripple characteristics.

## Materials and methods

Full experimental procedures are described elsewhere (Supplementary materials).

Animal experiments were performed with the approval of the Animal Experiment Ethics Committee at the University of Tokyo (approval number: A2025P007). Five 7-to 8-week-old ICR mice were used in this study. Ramelteon (R0216, Tokyo Chemical Industry, Japan) was dissolved in dimethyl sulfoxide and saline at a final concentration of 0.3 mg/ml.^11)^ The basic procedure of the stereotaxic surgery was conducted in accordance with our previous literature.^11)^ Briefly, one wire electrode was implanted into the trapezius to record EMGs. Nichrome wires were stereotaxically implanted into the PFC and dorsal hippocampus to record LFPs. Each drug (vehicle or ramelteon) was intraperitoneally administered to mice 20 min before recording. Mice were treated with vehicle and ramelteon in a crossover design.^6,15,16)^ The digitized biosignals were amplified, digitized at 2 kHz, and transferred to a data acquisition device (CerePlex Direct, Blackrock Neurotech). LFPs were recorded for 2 h in their home cages. Behavioral tasks were performed mostly during the nocturnal period for mice. After recordings, animals were anesthetized and transcardially perfused, followed by decapitation. The brains were soaked in 4% PFA overnight for post-fixation at 4°C. For cryoprotection, the brains were then immersed in 30% sucrose (in 0.01 M PBS in the tube) at 4°C until they sank to the bottom, which typically occurred within 24 h. Cryoprotected brains were coronally sectioned at 50 μm. Serial slices were processed for cresyl violet staining. The positions of all electrodes were confirmed by identifying tracks in the histological tissue (Fig. 1A, B). Data were excluded from the subsequent analysis if the electrode position was outside the target brain region.

**Figure 1.**
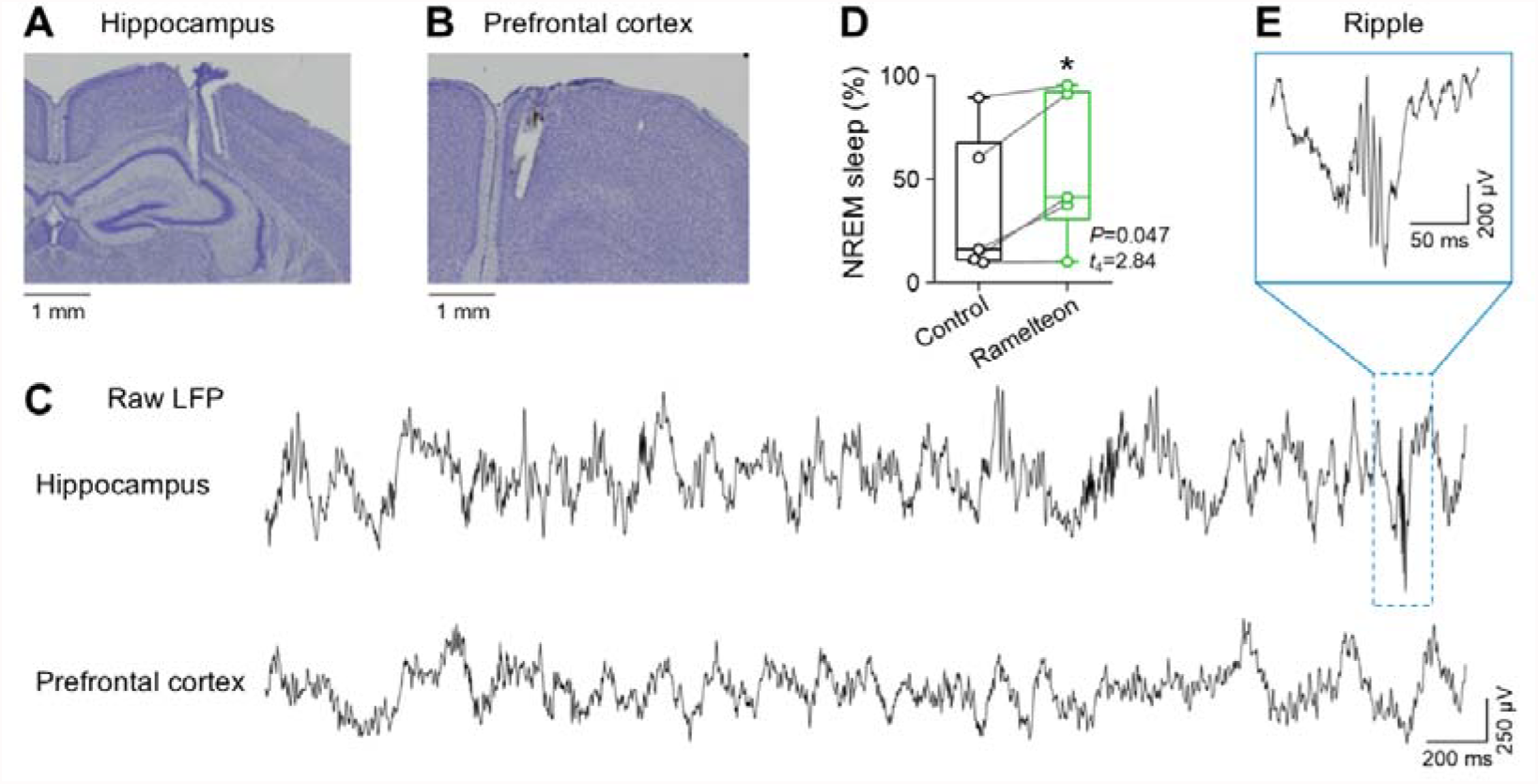
Ramelteon affects NREM sleep in mice. ***A***, A representative Nissl-stained section showing the electrode track in the hippocampus. ***B***, Same as *A*, but for the PFC. ***C***, Raw LFP traces simultaneously recorded from the hippocampus (*top*) and PFC (*bottom*). A representative ripple is indicated by *blue*. ***D***, Percentage of NREM sleep in control (*black*) and ramelteon-administered (*green*) groups. Ramelteon significantly increased the NREM sleep percentage (*P* = 0.047, *t*_4_ = 2.84, *n* = 5 mice, paired *t*-test). ***E***, A representative hippocampal ripple. The dashed box in *C* is magnified to show a sharp-wave ripple event recorded in the hippocampus. *Abbreviations*: NREM, non-rapid eye movement; PFC, prefrontal cortex; LFP, local field potential.

Data analysis was conducted using MATLAB. Summarized data were reported as the mean ± standard error of the mean. The significance level was set at 0.05. Vigilance states were classified based on EMG and LFP signals. Hippocampal LFPs were analyzed to detect ripples using a multi-stage protocol. For each detected ripple, several quantitative features were calculated to characterize its physiological properties: the amplitude, intra-ripple instantaneous frequency, duration, and asymmetry index.

## Results & Discussion

We performed chronic LFP recordings from the dorsal hippocampus and PFC in mice to investigate whether and how ramelteon affected hippocampal ripples during sleep (Fig. 1). Using neck EMGs and hippocampal LFPs (Fig. 1A–C), we evaluated how ramelteon treatment affected sleep architecture. Acute administration of ramelteon significantly increased the occupancy of NREM sleep during the 2-h recording period (37.3 ± 16.1% (control) *vs*. 55.0 ± 16.5% (ramelteon), *P* = 0.047, *t*_4_ = 2.84, *n* = 5 mice, paired *t*-test; Fig. 1D), consistent with our previous reports using rats.^6,16,17)^

We next examined the characteristics of hippocampal ripples during NREM sleep (Fig. 1C, E). We found that the occurrence rate of hippocampal ripples was significantly higher in the ramelteon-treated group than in the control group (0.021 ± 0.017 Hz (control) *vs*. 0.087 ± 0.069 Hz (ramelteon), *P* = 0.03, *n* = 5 mice, exact permutation test; Fig. 2A). Furthermore, the amplitude of ripples was significantly increased following ramelteon administration (175.9 ± 1.9 μV (control) *vs*. 278.9 ± 1.5 μV (ramelteon), *P* = 1.90 × 10^−163^, *D* = 0.64, *n* = 541 (control) and 2850 (ramelteon) events from 5 mice each, Kolmogorov-Smirnov test; Fig. 2B). We also observed a significant increase in the instantaneous frequency of the oscillations within ripples (152.4 ± 0.3 Hz (control) *vs*. 153.2 ± 0.2 Hz (ramelteon), *P* = 0.02, *D* = 0.07, *n* = 541 (control) and 2850 (ramelteon) events from 5 mice each, Kolmogorov-Smirnov test; Fig. 2C), but the low effect size (*i.e*., *D* = 0.07) indicated ramelteon treatment had a small effect on the ripple frequency. In contrast, other structural features of the ripples were not significantly affected. There were no significant differences in the ripple duration (34.2 ± 0.5 ms (control) *vs*. 33.7 ± 0.2 ms (ramelteon), *P* = 0.35, *D* = 0.04, *n* = 541 (control) and 2850 (ramelteon) events from 5 mice each, Kolmogorov-Smirnov test; Fig. 2D) or asymmetry index (−0.01 ± 0.01 (control) *vs*. −0.03 ± 0.01 (ramelteon), *P* = 0.07, *D* = 0.06, *n* = 541 (control) and 2850 (ramelteon) events from 5 mice each, Kolmogorov-Smirnov test; Fig. 2E).

**Figure 2.**
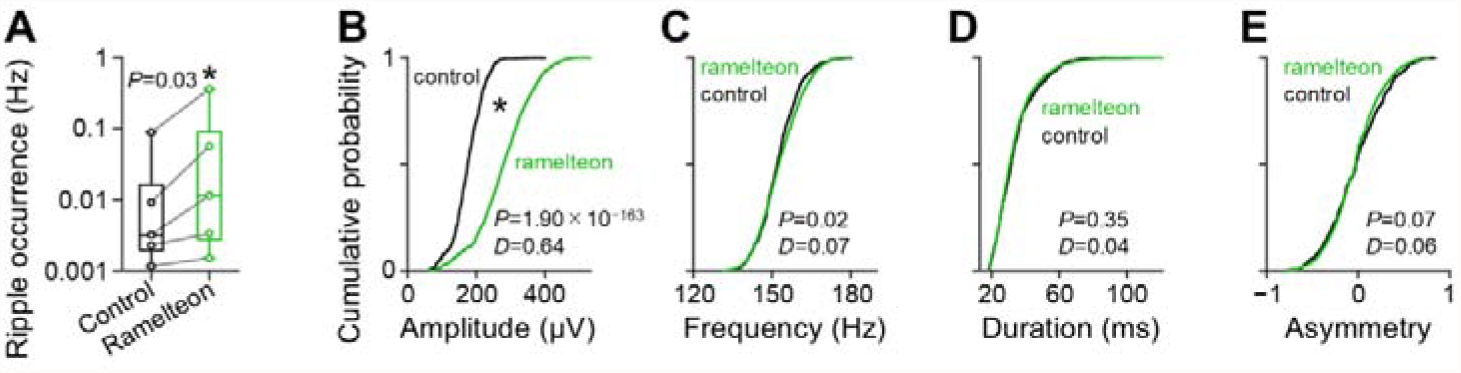
Characteristics of hippocampal ripples under ramelteon administration. ***A***, Ripple occurrence rate (Hz) in the control (*black*) and ramelteon (*green*) groups. Ramelteon significantly increased the frequency of ripple occurrence (*P* = 0.03, exact permutation test). ***B***, Cumulative probability distributions of ripple amplitude in control (*black*) and ramelteon (*green*) groups. Ramelteon significantly increased ripple amplitude (*P* = 1.90 ×10^−163^, *D* = 0.64, Kolmogorov-Smirnov test). ***C***, Same as *B*, but for intra-ripple instantaneous frequency. Ramelteon significantly increased ripple frequency, but the effect size was relatively small (*P* = 0.02, *D* = 0.07, Kolmogorov-Smirnov test). ***D***, Same as *B*, but for ripple duration. No significant difference was found between the two groups (*P* = 0.35, *D* = 0.04, Kolmogorov-Smirnov test). ***E***, Same as *B*, but for asymmetry index. No significant difference was found between the two groups (*P* = 0.07, *D* = 0.06, Kolmogorov-Smirnov test).

The increase in NREM sleep occupancy is consistent with the known pharmacological profile of ramelteon as an MT1/MT2 receptor agonist. Beyond its sleep-promoting effect,^18)^ our results reveal that ramelteon facilitates hippocampal ripple dynamics. The increased occurrence rate of ripples (Fig. 2A) suggests that ramelteon facilitates the transition of the hippocampal network into a ripple-generating state during sleep.

We also found a significant enhancement in ripple amplitude in ramelteon-treated mice (Fig. 2B). Since ripple amplitude is thought to reflect the extent of synchronous activity in pyramidal cells and their excitatory inputs,^20)^ our findings suggest that ramelteon enhances the synchronization of hippocampal neuronal ensembles during ripples. This modulation is likely mediated through melatonin receptors expressed in the hippocampus. Melatonin signaling has been believed to influence the excitation-inhibition balance by modulating GABAergic neurotransmission.^21,23)^ We speculate that ramelteon may shift the excitation-inhibition balance to favor the recruitment of larger population bursts within the CA1 network.

Ripples are essential for memory consolidation during sleep.^10,11)^ Consistently, we previously demonstrated that ramelteon improves object recognition memory in mice.^18)^ Combined with the current findings, it is plausible that the enhancement of ripple occurrence and amplitude during NREM sleep provides a potential neurophysiological basis for the procognitive properties of ramelteon. By refining the ripple-associated signals, ramelteon may facilitate more efficient information transfer from the hippocampus to downstream targets, such as the PFC, thereby promoting long-term memory consolidation. Our study identifies hippocampal ripples as a physiological target of ramelteon, offering new insights into how melatonin receptor agonists support cognitive function through the refinement of sleep-dependent neural oscillations.

## Acknowledgments

This work was supported by JST ERATO (JPMJER1801), JST Moonshot R&D (JPMJMS2012), the Institute for AI and Beyond of the University of Tokyo, JSPS KAKENHI (22K21353), AMED Brain/MINDS 2.0 (24wm0625207s0101; 24wm0625401h0001; 24wm0625502s0501) (to Y. Ikegaya), and JSPS KAKENHI (25K18705) (to N. Matsumoto).

