## Supplementary Methods for "Ramelteon facilitates hippocampal ripple occurrence and amplitude in mice"

### *Nakashima et al., Supplementary Materials*

#### Materials and methods (full)

##### Ethical approvals

Animal experiments were performed with the approval of the Animal Experiment Ethics Committee at the University of Tokyo (approval number: A2025P007) and according to the University of Tokyo guidelines for the care and use of laboratory animals. While our experimental protocols mandate humane euthanasia of animals if they exhibit any signs of pain, prominent lethargy, and discomfort, such symptoms were not observed in any of mice tested in this study. All efforts were made to minimize the animals' suffering.

##### Animals

Five 7- to 8-week-old ICR mice (Japan SLC, Japan) were housed in groups under conditions of controlled temperature and humidity (22 ± 1℃, 55 ± 5%) and maintained on a 12:12-h light/dark cycle (lights off from 7:00 a.m. to 7:00 p.m.) with *ad libitum* access to food and water. Mice were acclimated to an experimenter via daily handling for at least 3 days before experiments.

##### Drug

Ramelteon ($C_{16}H_{21}NO_2$; R0216, Tokyo Chemical Industry, Japan) was dissolved in dimethyl sulfoxide at a concentration of 30 mg/ml. This solution was diluted with saline solution to final concentrations of 0.3 mg/ml immediately before use. Saline with 1% dimethyl sulfoxide (*i.e.*, 0 mg/ml ramelteon) was used as the vehicle (control).

##### Surgery

A recording interface assembly was prepared as previously described. In short, the assembly was composed of an electrical interface board (EIB) (EIB-36-PTB, Neuralynx, USA) and shell and core bodies custom-made by a 3-D printer. The EIB had a sequence of metal holes for connections with wire electrodes. A given individual hole was conductively connected with one end of the insulated wire (~5 cm) using attachment gold pins, whereas the opposite end was soldered to a corresponding individual electrode during surgery.

The basic procedure of the stereotaxic surgery was conducted in accordance with our previous literature. General anesthesia was induced and maintained with 4% and 1–2% isoflurane gas, respectively, with careful inspection of the animal's condition during the whole surgical procedure. Veterinary ointment was applied to the mouse's eyes to prevent drying.

After complete anesthesia was confirmed, the electrodes for nuchal electromyograms (EMGs) were implanted as described previously. Briefly, the mouse was mounted onto a stereotaxic apparatus (SR-6M-HT, Narishige, Japan). One wire electrode (AS633, Cooner Wire, USA) was implanted into the trapezius to record EMGs. The mouse was then laid on its stomach and head-fixed in a stereotaxic apparatus using a nose clamp and bilateral ear bars. The scalp was then removed with a surgical knife. Circular craniotomies with a diameter of approximately 1.0 mm were performed using a high-speed dental drill (SD-102, Narishige). Penetrative nichrome wires with a bare diameter of 50.8 μm (762000, A-M Systems, USA) were stereotaxically implanted unilaterally into the PFC (1.6 mm anterior and 0.5 mm lateral to bregma) and dorsal hippocampus (2.0 mm posterior and 2.0 mm lateral to bregma) to record LFPs. Two stainless-steel screws were additionally implanted into the bone above the cerebellum (6.0 mm posterior and 1.5–2.0 mm bilateral to bregma) as ground and reference electrodes. Each of the open edges of ground and reference electrodes was soldered to the corresponding open edge of insulated wires of the recording interface assembly. This assembly including all electrodes was secured to the skull using dental cement.

Following surgery, each mouse was allowed to recover from anesthesia and was housed individually with free access to water and food. For the first 3–5 d after surgery, the health condition of animals was carefully checked every day.

##### In vivo electrophysiology

Each home cage was surrounded by white corrugated plastic sheets to prevent a mouse from escaping during recording. Mice were habituated to the recording interface assembly for 15 min (per day) for 2 d.

Each drug (vehicle or ramelteon (3.0 mg/kg)) was intraperitoneally administered to mice 20 min before recording. Mice were treated with vehicle and ramelteon in a crossover design, in which the interval between administration of the two drugs was set at over 36 h and the administration order of the drugs was randomly predetermined.

#### Materials and methods (full)

(continued from the previous page)
The EIB of the recording interface assembly was connected to digital headstage (CerePlex μ, Blackrock Neurotech, USA). The digitized signals were amplified and transferred to a data acquisition device (CerePlex Direct, Blackrock Neurotech) via interface cables. Electrophysiological signals were digitized at a sampling rate of 2 kHz. LFPs were recorded for approximately 2 h in their home cages (W190 × D260 × H128 mm; MC-2K, ITECH, Japan). A web camera (MCM-300A, Gazo, Japan) was placed above the apparatus to monitor the animals' behavior at 60 fps. Camera's strobe signals were transmitted to the data acquisition system (CerePlex Direct) to synchronize with electrophysiological signals using a custom-made application. Behavioral tasks were performed mostly during the nocturnal period for mice (*i.e.*, 7 a.m. to 7 p.m.).

**Histology**
After the recordings, animals were deeply anesthetized by intraperitoneal injection of an overdose of urethane and transcardially perfused with 0.01 M phosphate-buffered saline (PBS; pH 7.4) and 4% paraformaldehyde (PFA) in 0.01 M PBS, followed by decapitation. The brains were soaked in 4% PFA overnight for post-fixation at 4℃. For cryoprotection, the brains were then immersed in 30% sucrose (in 0.01 M PBS in the tube) at 4℃ until they sank to the bottom, which typically occurred within 24 h. The brains were coronally sectioned at a thickness of 50 μm using a microtome (Leica SM2010 R, Leica Biosystems, Germany). Serial slices were mounted on glass slides using 0.2% gelatin (in 0.01 M PBS) and processed for cresyl violet staining. For cresyl staining, the slices were rinsed in water, ethanol, and xylene, counterstained with cresyl violet, and coverslipped with malinol (2009-1, Muto Pure Chemicals, Japan). The positions of all electrodes were confirmed by identifying tracks in the histological tissue (Fig. 1A, B). Data were excluded from the subsequent analysis if the electrode position was outside the target brain region. Cresyl violet-stained images were acquired using a phase-contrast microscope (BZ-X810, Keyence, Japan).

**Data analysis**
**Statistics**
Data analysis was conducted using MATLAB. Summarized data were reported as the mean ± standard error of the mean. The significance level was set at 0.05. For pairwise comparisons, the normality of the differences between paired datasets was evaluated by the Shapiro-Wilk test. Paired *t*-tests were employed when the differences followed a normal distribution; otherwise, an exact paired permutation test was applied, based on all $2^n$ possible sign-flips of the differences ($n$ = 5 mice).

**Vigilance state scoring**
The classification of vigilance states was carried out using a sliding window approach with a window size of 2 s for feature extraction. Raw EMG and LFP signals were first preprocessed by trimming to remove noise. Specifically, signal amplitudes were clipped at the 99.5$^{th}$ percentile to minimize the influence of extreme artifacts. The trimmed EMG signals were bandpass-filtered between 20–200 Hz to extract muscle activity.
For movement detection, the root-mean-square of the filtered EMGs (called $EMG_{RMS}$, hereafter) was calculated for each 2-s time window. The threshold for movement (*i.e.*, $Th_{EMG\_mov}$) was set at $\mu_{RMS\_10} + k \times \sigma_{RMS}$, where $\mu_{RMS\_10}$ was the mean of the $EMG_{RMS}$ values below the 10$^{th}$ percentile and $\sigma_{RMS}$ was the standard deviation of the $EMG_{RMS}$ values. The scaling factor $k$ was empirically optimized for each video recording by visually cross-validating the detected movement segments with the recorded behavior in the synchronized video clips. A given window $t$ was classified as behaving (active awake) if the $EMG_{RMS}$ values exceeded $Th_{EMG\_mov}$. This binary classification was then upsampled to the original sampling rate by repeating each window value (*e.g.*, 0 for $EMG_{RMS}(t) < Th_{EMG\_mov}$ and 1 for $EMG_{RMS}(t) > Th_{EMG\_mov}$). To refine the state detection, two postprocessing steps were applied: first, short bouts of movement lasting less than 5 s were removed; second, gaps between movement periods shorter than 20 s were merged to ensure temporal continuity. Subsequently, for each 2-s window, the delta power (1–4 Hz) in the PFC and hippocampus and the theta (4–12 Hz) power in the hippocampus was calculated. The hippocampal theta-to-delta ratio (*i.e.*, θ/δ) was determined to distinguish between REM and NREM sleep.

#### Materials and methods (full)

(continued from the previous page)
To objectively determine the thresholds for REM/NREM sleep state scoring, Otsu's method was applied to the distribution of the extracted neural features. The hippocampal θ/δ and PFC delta power were first base-10 log-transformed ($\log_{10}$) to highlight their (bimodal) characteristics. The optimal thresholds for distinguishing REM sleep (based on the hippocampal θ/δ ($HPC_{\theta/\delta}$)) and NREM sleep (based on the PFC delta power ($PFC_{delta}$)) were then automatically calculated by maximizing the inter-class variance of the respective histograms and called as $Th_{HPC\_\theta/\delta}$ and $Th_{PFC\_delta}$, respectively.

Based on these thresholds, each 2-s window was assigned to one of four vigilance states: active awake (previously identified as behaving by the movement detection described above), quiet awake, NREM sleep, or REM sleep. For the remaining non-behaving windows, a low-EMG reference threshold ($Th_{EMG\_low}$) was determined by averaging the EMG RMS values below the 0.5th percentile to identify periods of muscle atonia. A given 2-s window *t* was classified as REM sleep if they exhibited both low EMG activity ($EMG_{RMS}(t) < Th_{EMG\_low}$) and a high hippocampal θ/δ ($HPC_{\theta/\delta}(t) > Th_{HPC\_\theta/\delta}$). Among the remaining windows, those with high PFC delta power ($PFC_{delta}(t) > Th_{PFC\_delta}$) were categorized as NREM sleep, while the rest were labeled as quiet awake. Finally, these bin-wise state labels were upsampled to the original sampling rate by repeating each label value for the duration of the window, with the final sequence adjusted to match the total number of recording sample points.

**Hippocampal ripple detection**
Hippocampal LFPs were analyzed to detect ripples using a multi-stage protocol. Raw LFP signals were, where necessary, first resampled to a consistent sampling rate of 2,000 Hz. Ripple detection was performed by bandpass-filtering the signal between 120 and 200 Hz using a third-order Butterworth filter.[5)] The power envelope was then calculated by squaring the filtered signal and applying a 5-ms moving average window. This envelope was Z-score normalized to generate a normalized squared signal (NSS).
Event identification employed a double-thresholding approach where an onset and offset were recognized when the NSS exceeded 2 × standard deviations (SD), and the peak power within the segment was required to exceed 5 × SD. Neighboring events with an interevent interval of less than 15 ms were merged into a single event. To ensure physiological validity, only events with a duration between 15 and 250 ms were retained.

To further exclude artifacts, each candidate underwent stringent validation based on multiple criteria. Events were rejected if the peak filtered amplitude exceeded 700 μV or if the raw LFP amplitude exceeded 1,000 μV. Using the Hilbert transform, the instantaneous frequency and phase were calculated to ensure that each event exhibited at least three continuous cycles of oscillation within the 120–200 Hz range. Furthermore, a peakiness index was calculated as the ratio of peak amplitude to the mean envelope amplitude, and events with an index greater than 3 were excluded as spiky artifacts. Subsequently, the remaining ripple candidates were subjected to manual visual inspection by three skilled experimenters. Artifacts identified during this manual curation were excluded from the final dataset to ensure high-fidelity detection of hippocampal ripples.

For each detected ripple, several quantitative features were calculated to characterize its physiological properties. The temporal boundaries were defined by the onset ($t_{onset}$) and offset ($t_{offset}$), representing the time points where the NSS first exceeded and subsequently fell below the edge threshold of 2 × SD. Within this interval, the peak time ($t_{peak}$) was identified as the time point at which the NSS reached its maximum value. At this specific peak time, the instantaneous frequency was determined by calculating the derivative of the smoothed phase obtained via the Hilbert transform. The maximum amplitude was defined as the peak of the signal envelope, calculated as the instantaneous magnitude of the complex-valued signal derived from the Hilbert transform. To evaluate the temporal structure of the waveform, the asymmetry index was computed using these three time points (*i.e.*, $t_{onset}$, $t_{peak}$, and $t_{offset}$) as $((t_{peak} - t_{onset}) - (t_{offset} - t_{peak})) / (t_{offset} - t_{onset})$.[32)] This formula calculated the difference between the duration of the rising phase ($t_{peak} - t_{onset}$) and the falling phase ($t_{offset} - t_{peak}$), normalized by the total duration of the event. The asymmetry index ranged from −1 to 1, where a value of 0 indicates a symmetric envelope. A positive index indicates that the peak power is shifted toward the end of the ripple (offset), whereas a negative index indicates a shift toward the beginning (onset).
